# Mechanistic Insights into HIV-1 Capsid Interactions with CPSF6

**DOI:** 10.64898/2026.09.05.749640

**Authors:** Arpa Hudait, Kuntal Ghosh, Gregory A. Voth

**Affiliations:** Department of Chemistry, Chicago Center for Theoretical Chemistry, Institute for Biophysical Dynamics, and James Franck Institute, The University of Chicago, Chicago, IL 60637, USA

**Keywords:** HIV-1 capsid, virus-host interactions, nuclear entry, capsid curvature, molecular dynamics, coarse-grained

## Abstract

A crucial stage of the HIV-1 life cycle is the docking of the viral capsid at, and its translocation through, the nuclear pore complex (NPC). During this process, the capsid interacts with a series of cellular host factors that regulate efficient nuclear entry. One such host factor is cleavage and polyadenylation specificity factor 6 (CPSF6), which plays a critical role in efficient HIV-1 nuclear entry and integration. Experimental evidence suggests that nucleoporins (NUPs), such as NUP153, initially engage the capsid, followed by CPSF6 binding and oligomerization during subsequent stages of nuclear import. However, the mechanistic basis by which CPSF6 binds to the capsid and subsequently oligomerizes remains poorly understood. To address this question, we have developed bottom-up coarse-grained (CG) models that explicitly simulate the process of CPSF6 binding and oligomerization on the HIV-1 capsid. Our simulations show that CPSF6 assembles into a mesh-like coating surrounding the capsid, consistent with experimental observations. Furthermore, we then demonstrate that CPSF6 binding is dependent on the intrinsic curvature of the capsid lattice. Finally, we investigate how perturbing CPSF6-CPSF6 interactions alters binding, oligomerization dynamics, and curvature dependence. Together, these findings provide new mechanistic insights into how CPSF6 regulates HIV-1 nuclear entry.

## INTRODUCTION

After Human immunodeficiency virus type 1 (HIV-1) enters the cell, several distinct steps guide the mature viral capsid containing genomic material into the nucleus [1,2]. The interior of the capsid contains the viral genome and enzymes and is the site of reverse transcription [3,4]. Uncoating of the capsid releases the genome to allow integration [5]. The nuclear pore complex (NPC) is a multiprotein assembly embedded in the nuclear envelope that regulates the passage of the HIV-1 capsid [6,7]. Multiple cellular host factors at the cytoplasmic side, NPC central channel, and nuclear basket facilitate the passage of HIV-1 capsid from the cytoplasmic side of the NPC to the nucleus [8-11]. The sequential steps that regulate the nuclear entry of the capsid are potential targets for capsid (CA) inhibitor molecules [12,13].

Cellular host factor proteins cleavage and polyadenylation specificity factor 6 (CPSF6) and nucleoporin NUP153 interact with the viral capsid at various stages of nuclear entry. NUP153, present in high concentrations at the nuclear basket, mediates translocation of the capsid from the NPC central channel to the nuclear end [14,15]. At the nuclear basket, CPSF6 binds to CA protein hexamers, mediates the handover of the capsid from NUP153, transports the capsid to the interior of the nucleus, and promotes integration [16-19]. Experiments have suggested that abrogating CPSF6 binding to CA through depletion of CPSF6 or mutation of CPSF6 binding CA residues stops the capsid at the NPC [16,20]. Therefore, the interaction of CPSF6 and CA appears to be essential for the efficient handover of the capsid from the nuclear basket to the nucleoplasm.

CPSF6 and NUP153 are intrinsically disordered and contain a phenylalanine-glycine (FG)- containing peptide sequence (CPSF6_313–327_ and NUP153_1409–1423_). CPSF6 and NUP153 can also oligomerize to higher-order complexes and hydrogels [21-23]. FG peptide of the cellular host factor proteins is flanked by disordered prion-like low complexity regions (LCRs). Prion-like regions typically are rich in uncharged amino acids and are known to induce oligomerization and liquidliquid phase separation [24,25]. A recent combined cryo-EM structural, virology, and biochemical analysis study has discovered that CPSF6 self-assembly is mediated by LCR-LCR interactions [26]. CPSF6 FG-motif exhibits low affinity binding to the CA hydrophobic pocket. Therefore, beyond the interaction of CPSF6 FG peptide with CA, LCR-LCR interactions provide CPSF6 binding avidity to the capsid lattice for efficient virus-host interactions. However, to the best of our knowledge, the mechanistic basis of these LCR-LCR interactions that drive capsid entry, is not well understood.

In this work, we performed large-scale coarse-grained (CG) molecular dynamics (MD) simulations to elucidate the dynamic process of CPSF6 oligomerization and its concurrent binding to the HIV-1 capsid. Our simulations suggest the molecular mechanism by which CPSF6 progressively coats the capsid, from the initial binding of monomers to the assembly of higherorder oligomers. We further performed extensive analyses of the effects of LCR-LCR interactions on CPSF6 binding dynamics, demonstrating that CPSF6 binding is dependent on the intrinsic curvature of the capsid lattice. Taken together, our results provide mechanistic insights into the molecular interactions governing CPSF6-mediated regulation of HIV-1 nuclear entry.

## RESULTS

### Coarse-grained modeling and simulations

Mature HIV-1 capsid cones typically consist of ∼1500 CA monomers [27]. Therefore, simulating the full capsid in addition to other components relevant to nuclear entry is computationally infeasible. CG MD simulations are particularly efficient in accessing length and time scales far beyond the reach of simulations performed with fully atomistic detail. CG MD has allowed us to access highly complex and relatively slow processes involving viral dynamics, such as HIV-1 capsid lattice assembly and uncoating [28-31], viral restriction factor assembly [32], immature Gag assembly [33,34], formation of ESCRT complexes at the budding sites [35] and envelope glycoprotein oligomerization [36]. To additionally help us to surmount the computational cost, we employ “solvent-free” CG molecular models (**Fig. 1**). This phrase signifies that the net effects of solvent are folded in the nature of the CG interactions, thereby reducing the overall number of CG particles (aka “beads”) in the system and decreasing the overall computational cost [37-41].

**FIGURE 1.**
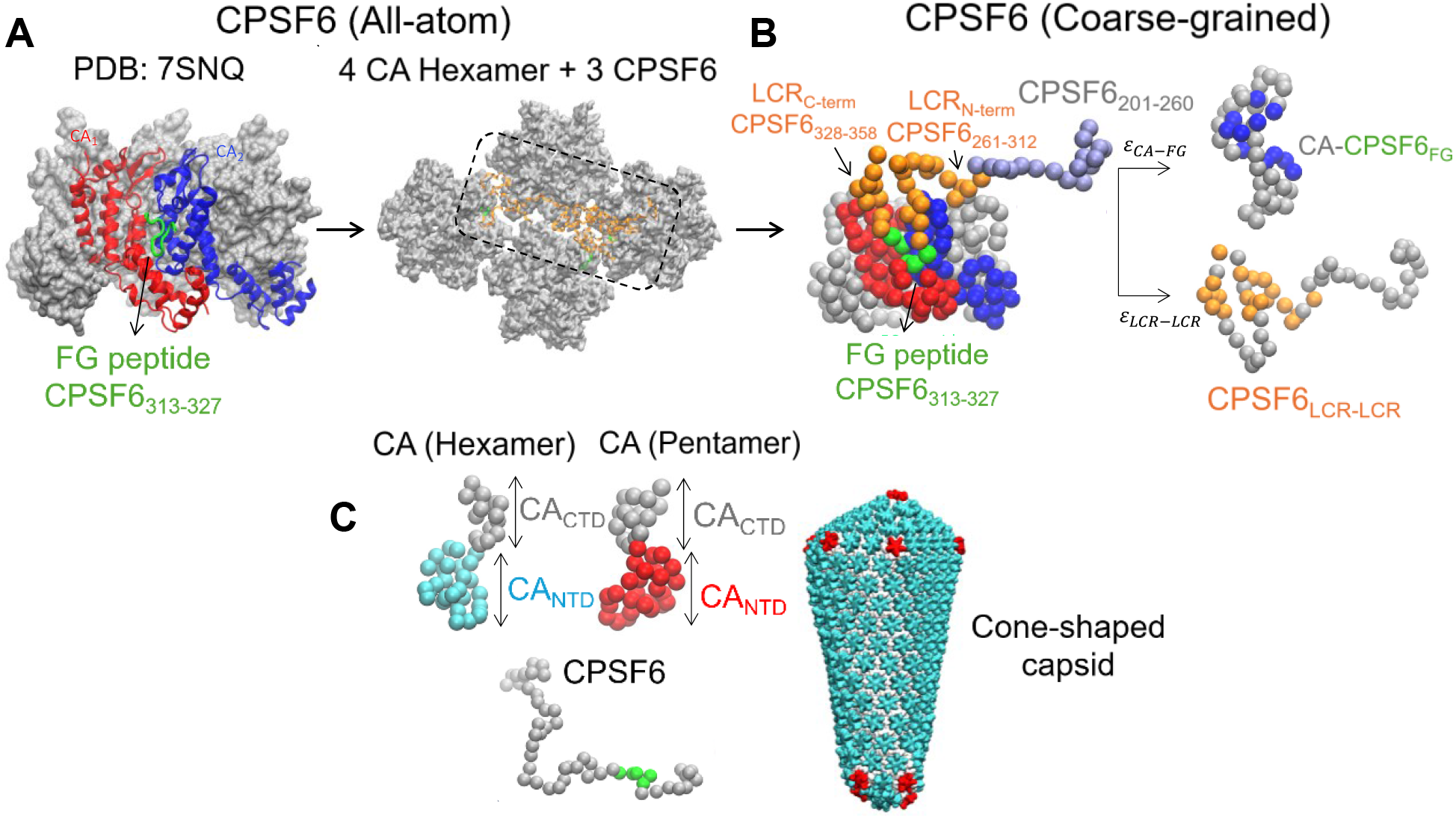
CG model of CPSF6, CA and the associated molecular interactions. **(*A*)** The X-ray crystal structure of FG peptide (CPSF6_313-327_) bound to CA hexamer (PDB: 7SNQ) is shown in the left panel. The FG peptide is shown in green ribbon. The CA monomers contiguous to the FG peptide are shown in blue and red ribbons. The rest of the CA monomers are shown in silver surface representation. The reference all-atom simulation of the 4 CA hexamer and 3 GSTCPSF6_261-358_ complex from which the CG molecular interactions are derived are shown in the right panel. The initial configuration of the system is constructed from the CPSF6 assembly templated by CA nanotube (EMD-27619). The dotted region shows the 3 interacting CPSF6 chains in orange ribbons. The CA hexamers are shown in silver. **(*B*)** CG representation of a CPSF6 chain bound to CA hexamer. The adjoining CA monomers that create the hydrophobic pocket for FG peptide binding are shown in red and blue spheres. The FG peptide is shown in green spheres. The N-terminal LCR (CPSF6_261-312_) and C-terminal LCR (CPSF6_328-358_) segments are shown in orange spheres. The N-terminal non-LCR segment (CPSF6_201-260_) is shown in ice blue spheres. The upper right panel shows (as blue spheres) the CG sites of a CA monomer involved in associative interactions with the CG CPSF6 FG peptide. The other CG sites of CA (not involved in associative interactions with the FG peptide) are shown in silver spheres. The lower right panel shows (as orange spheres) the CG sites of a CPSF6 monomer involved in LCR-LCR associative interactions. The other CG sites (not involved in associative LCR-LCR interactions) are shown in silver spheres. **(*C*)** CG representation of the CA monomer is shown in the upper left panel. The CA_NTD_ domains are shown in cyan and red for the hexamer and pentamers, respectively. CA_CTD_ domain of both hexamer and pentamers are shown in silver spheres. The right panel shows the cone-shaped capsid. In our CG model, CPSF6 FG peptide selectively binds to CA monomers of the hexamers.

The “bottom-up” CG molecular models [42] of CPSF6, CA-CPSF6, CPSF6-CPSF6 interactions were derived from previously reported all-atom (AA) MD simulations [26]. The AA MD simulations consisted of 3 copies of CPSF6_261-358_ bound to 4 CA hexamers. The CG CPSF6 monomer is modeled as a linear polymer chain with a resolution of 3 residues for each CG site. In addition to CPSF6_261-358_, we added a 20-bead polymer chain to the N-terminal end to model CPSF6_201-260,_ which does not have any experimentally characterized LCR-like interactions and, therefore, is oligomerization-incompetent. The resulting CPSF6_200-358_ CG model consists of 53 CG sites (**Fig. 1*A* and 1*B***). Each CA monomer consists of 46 CG sites, with an average CG model resolution of 5 residues per CG site, and CA-CA attractive interactions were used to model the entire mature capsid structures [11]. The CG-mapped trajectory of the atomistic CA-CPSF6 complex was used to derive the CG non-bonded associative interactions between CA-FG (*ε*_*CA*−*FG*_) and LCR-LCR (*ε*_*LCR*−*LCR*_). Complete details of the CG model are provided in *SI Methods*.

Using the aforementioned models, we extensively simulated the CPSF6 assembly templated by the cone-shaped capsid. In these simulations, we prepared a system with unbound CPSF6 distributed away from the capsid. The initial concentration of unbound CPSF6 is ∼ 2.9 CPSF6 proteins per CA hexamer in the simulations. Note, that there are 209 CA hexamers in the cone. In experimental structures, the mean binding stoichiometry of CPSF6 to CA hexamer is ∼2.4 [26]. Additional details of the simulation setup are provided in the *Methods*.

### Assembly of CPSF6 oligomers templated by HIV-1 capsid is driven by interchain LCR-LCR associative interactions

Structural characterization of CPSF6 bound to the IP6-stabilized CA nanotubes revealed CPSF6 targets CA hexamers through FG peptide association at a hydrophobic binding pocket [26]. Other host factors rich in FG repeats (such as NUP153) also bind to the same hydrophobic pocket [15]. The structurally disordered LCR segments of CPSF6 occupy the region between CA hexamers forming a continuous network of interacting chains extending outwards from the CA lattice. Additional virology experiments demonstrated that chimeric CPSF6 with non-LCR segments do not engage with HIV-1 cores. Here we used CG MD simulations to better elucidate the factors that drive oligomerization of CPSF6 templated by the HIV-1 capsid. The simulations performed here mimic events when the capsid passes through the NPC central channel and localizes at the nuclear basket. At this point, CPSF6 enrichment at the periphery of the basket and subsequent oligomerization coating the capsid facilitate the release of the capsid from the NPC to the nuclear interior [1,16] (**Fig. S1**).

First, we performed CG MD simulations of CPSF6 oligomerization using the wildtype (WT) LCR-LCR associative interaction strength derived from AA MD simulations (*ε*_*LCR*−*LCR*_ = 2.8 kcal/mol). Our CG MD simulations illustrate both CPSF6 binding and oligomerization progress simultaneously, indicating that sub-stoichiometric binding of CPSF6 to the capsid is sufficient to trigger extended oligomerization (**Fig. 2*A***). Time-series profiles in **Fig. 3** show the total number of CPSF6 bound (*N*_*bound*_) and number of oligomers (*N*_*oligo*_). Our previous all-atom (AA) MD simulations had demonstrated that a trimeric CPSF6 oligomer is a minimal self-assembling motif mediated by interchain LCR-LCR interactions [26]. Therefore, to characterize the stepwise CPSF6 nucleation dynamics, we calculated the time-series of trimeric CPSF6 clusters (*N*_*trimer*_) in the CG MD simulations (**Fig. S2**). We find that *N*_*trimer*_ increases up to 1300 × 10^6^ CG time steps and then decreases while the number of CPSF6 oligomers continues to grow. The time up to 1300 × 10^6^ CG time steps can be classified as the nucleation phase during which several trimeric motifs are formed. After that, the trimeric motifs recruit additional CPSF6 monomers forming extended oligomers. CPSF6 monomers can bind to unoccupied CA monomers driven by CA-FG interactions. However, there is a higher energetic driving force regulated by LCR-LCR associative interactions for an unbound CPSF6 monomer to attach to a preexisting trimeric or higher-order oligomer. Therefore, the LCR-LCR interactions provide additional driving force for CPSF6 binding to the capsid. We next determine the importance of LCR-LCR associative interactions for CPSF6 binding to the capsid. It must be noted that CG time is not a real time: it is a greatly accelerated time due to the CG modeling.

**FIGURE 2.**
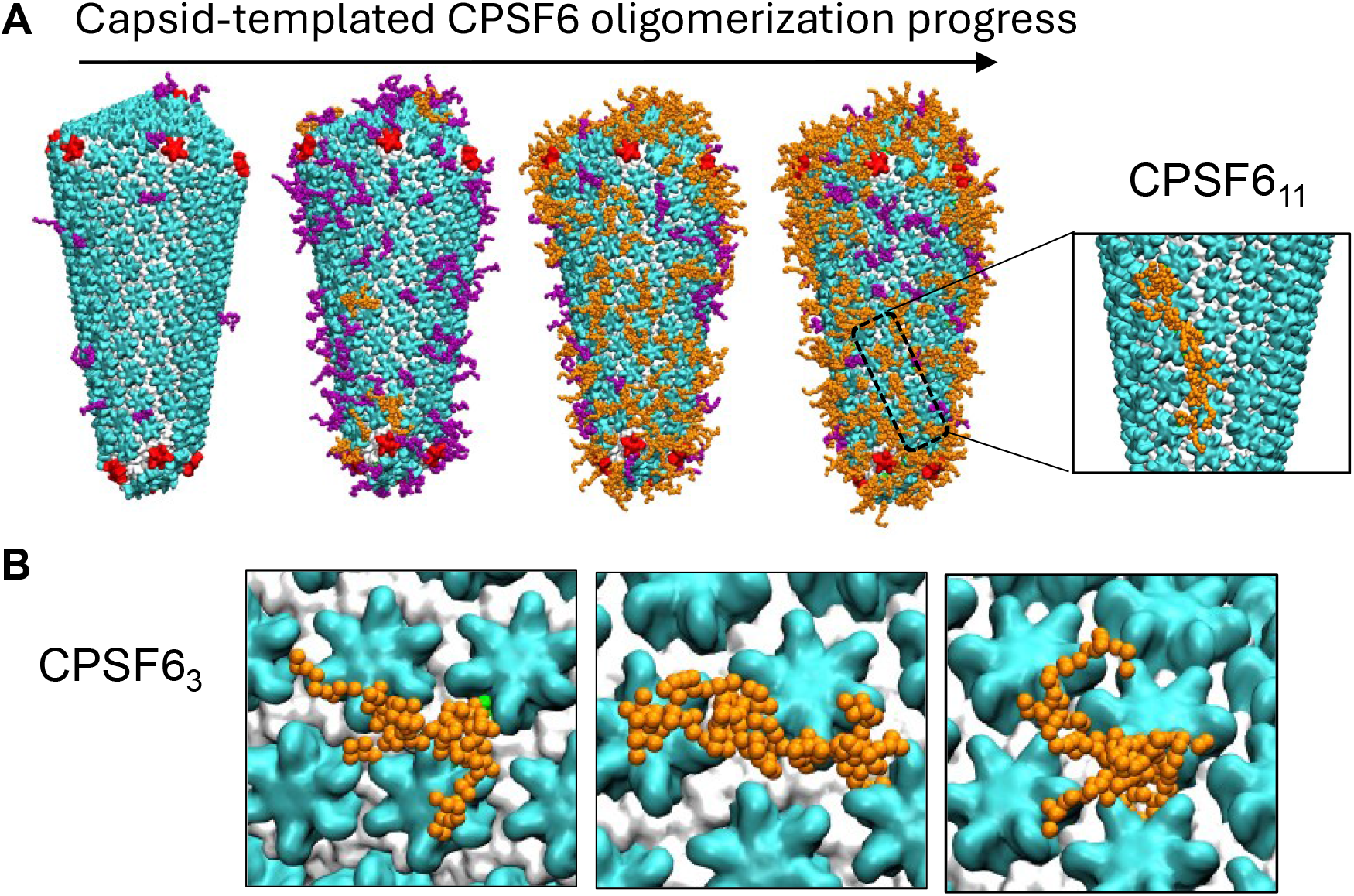
CPSF6 oligomerization templated by the cone-shaped capsid. **(*A*)** The snapshots depict CPSF6 oligomerization progress (from left to right) with simulation time. In the snapshots, the CA_NTD_ domain of capsid hexamers and pentamers are shown in cyan and red, respectively. The CA_CTD_ domain for both hexamers and pentamers is shown in silver. In all snapshots, only the FG peptide and flanking LCRs of CPSF6 are shown for visual clarity. The capsid bound CPSF6 monomers or dimers are shown in purple. The oligomerized clusters (*N*_*monomer*_ ≥ 3) are shown in orange. A zoomed-out snapshot of an oligomer containing 11 CPSF6 chains (CPSF6_11_) is shown. **(*B*)** Snapshots of different CPSF6 trimeric motifs (CPSF6_3_) bound to the capsid lattice.

**FIGURE 3.**
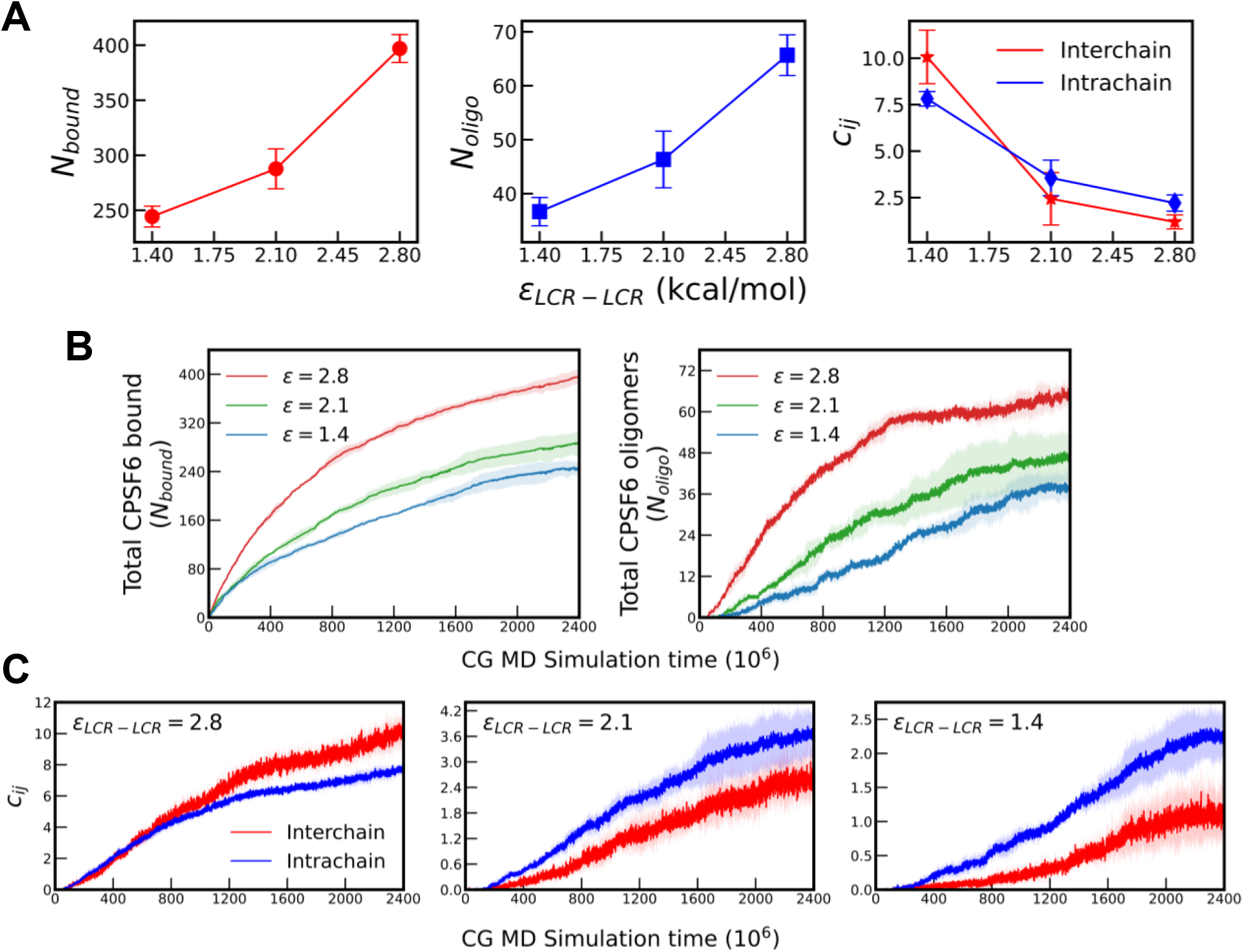
Role of LCR-LCR interactions (*ε*_*LCR*−*LCR*_ or simply *ε*) in modulating CA-CPSF6 interactions and CPSF6 oligomerization. **(*A*)** The number of total CPSF6 bound to the capsid lattice (*N*_*bound*_), total oligomerized clusters (*N*_*oligo*_), and cumulative inter and intrachain LCR-LCR contact parameter (*c*_*ij*_) for *ε*_*LCR*−*LCR*_ = 2.8 (WT), 2.1 and 1.4 kcal/mol. The mean and standard deviation values are calculated from the endpoint of the simulation trajectories for 3 replicas. ***(B)*** Time evolution of *N*_*bound*_ and *N*_*oligo*_ for different values of LCR-LCR interactions. ***(C)*** Time evolution of both intra and interchains *c*_*ij*_ for different values of LCR-LCR interactions.

We performed simulations by diminishing the value of *ε*_*LCR*−*LCR*_ of the WT CPSF6 LCRLCR interactions by 25% (*ε*_*LCR*−*LCR*_ = 2.1 kcal/mol) and 50% (*ε*_*LCR*−*LCR*_ = 1.4 kcal/mol) to create computationally mutated CPSF6 models. **Figure 3** shows the *N*_*bound*_ and *N*_*oligo*_ values of the mutated CPSF6 compared to the WT CPSF6 at the endpoint of the simulation trajectories. As the LCR-LCR interactions are diminished, there is a significant decrease in the oligomerization efficacy. Importantly, our simulations show that the progressive weakening of the LCR-LCR interactions impacts the degree of CPSF6 binding and hence illustrate the role of LCR-LCR interactions on the binding avidity. This is further demonstrated in another CPSF6 mutant; when the LCR interactions are completely turned off there is a 4-fold decrease in *N*_*bound*_ related to the WT CPSF6 (**Fig. 3*A*** and **Fig. S3**). To characterize the molecular details of LCR-mediated CPSF6 oligomerization, we calculated the LCR-LCR contact parameter (*c*_*ij*_) in the oligomerized clusters for the WT and mutated CPSF6 proteins (**Fig. 3*A*, Fig. 3*C*** and **Fig. S4**). The LCR-LCR contact parameter (*c*_*ij*_) is decomposed into intra- (*c*_*ij,intra*_) and interchain (*c*_*ij,inter*_) components (**Fig. 3*A***). For the WT CPSF6, we observe a higher degree of interchain contacts compared to intrachain contacts, in contrast to that for the mutated CPSF6 proteins. In other words, when bound to the capsid in the monomeric form, LCR segments can form multiple intrachain contacts due to the confined environment of the lattice. When another CPSF6 monomer binds in the vicinity, the driving force to establish interchain LCR-LCR contacts significantly diminishes for weaker LCRLCR interaction impeding oligomerization.

### CPSF6 binding to CA lattice has a curvature dependence

We next analyzed the dependence of CPSF6 binding on the curvature of the capsid lattice. As described in the aforementioned sections, we probe the binding dynamics of CPSF6 on a cone-shaped lattice, which is characterized by regions of varying curvature. In our simulations, we examine the mean local tilt angles of six nearest neighbor hexamers with respect to the hexamer bound to a CPSF6 moiety (**Fig. S5**). A lower local hexamer tilt angle indicates a flatter hexamer patch, whereas a higher local hexamer tilt angle indicates a region of higher curvature. We then compare the distribution of local tilt angles for hexamers hosting different numbers of CPSF6 oligomers. Our CG MD simulations reveal (**Fig. 4**) that the number of CPSF6 oligomers hosted by a hexamer has a clear dependence on the local curvature: hexamers hosting two oligomers (n = 2) peak at higher tilt angles than hexamers hosting a single oligomer (n = 1), and both are shifted relative to hexamers bound to CPSF6 monomers only (n = 0) suggesting that regions of higher curvature preferentially co-localize multiple CPSF6 oligomers, particularly for interaction strengths *ε*_*LCR*−*LCR*_ = 2.8 and 2.1 kcal/mol (**Fig. 4*A*** and **Fig. 4*B***). Hexamers hosting a single CPSF6 oligomer (n = 1) show a more modest extension into the higher-tilt regime. At *ε*_*LCR*−*LCR*_=1.4 kcal/mol this trend is weaker: the n = 2 distribution becomes dominated by its low-tilt mode overlapping the n = 0 population, with only a small secondary population at higher tilt (**Fig. 4*C***). Thus, the mode of curvature-dependent CPSF6 binding also depends on the LCR-LCR interactions. It is to be noted that n = 2 denotes hexamers to which two mutually disconnected CPSF6 oligomers are assigned by the plurality of their member chains being bound to that hexamer.

**FIGURE 4.**
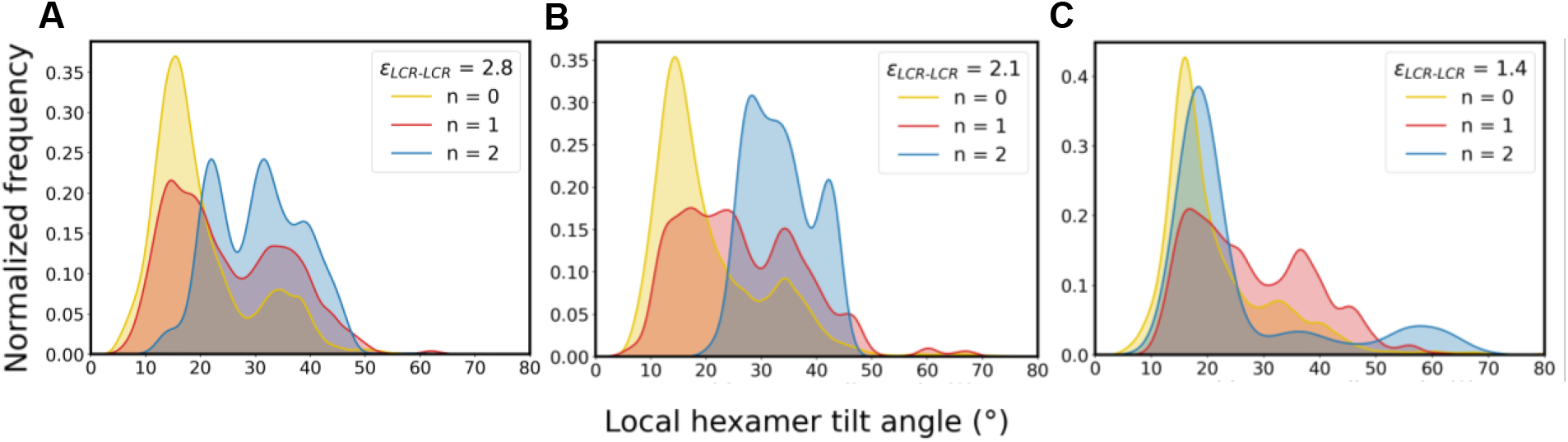
CPSF6 binding dependence on capsid curvature for three values of LCR-LCR interactions. **(*A*)** *ε*_*LCR*−*LCR*_= 2.8 **(*B*)** *ε*_*LCR*−*LCR*_= 2.1 **(*C*)** *ε*_*LCR*−*LCR*_= 1.4 kcal/mol. In all panels, n=0 denotes CA hexamers to which no oligomer is assigned. n=1 denotes hexamers hosting a single CPSF6 oligomer, and n=2 denotes hexamers hosting two CPSF6 oligomers.

## DISCUSSION AND CONCLUSIONS

In this work, we have systematically investigated the dynamic binding of CPSF6 oligomers to the HIV-1 capsid using bottom-up CG methodology. We probed the mechanism by which CPSF6 units interact with the hydrophobic binding pockets in the capsid, followed by CPSF6 oligomerization, as mediated by LCR-LCR interactions.

We furthermore identified the critical mechanistic features of capsid-mediated oligomerization of the cellular host factor CPSF6. We first developed a CG CPSF6 model, in which we account for the dominant contacts between LCR segments between adjoining CPSF6 chains bound CA hexamers from AA MD and express the LCR associative interactions through strictly pairwise interactions. Our simulations demonstrate that diminishing the LCR-LCR interactions also adversely impacts the oligomerization and binding efficiency of mutated CPSF6. Therefore, our simulations provide an underlying molecular-scale picture of the multivalent oligomerization mechanism of CPSF6 templated by the HIV-1 capsid. These results present the following mechanistic pathway for CPSF6 oligomerization. The low-affinity binding of the FG peptide allows the binding of multiple CPSF6 monomers throughout the capsid lattice. The FG peptide occupies the hydrophobic CA binding pocket, and the intrinsically disordered LCR condensates extend away from the capsid surface. The conformational flexibility of LCRs is advantageous to establish extensive LCR-LCR contacts between an incoming CPSF6 monomer and the already bound CPSF6 oligomers. This is unlike self-assembly between structured proteins, where the orientation of the approach is key to successful oligomer formation [28,33]. As our simulations demonstrate, energetically favorable LCR-LCR interactions and oligomerization also distinctly augment host factors binding to the capsid lattice.

In addition, we show that CPSF6 binding depends on the curvature of the capsid lattice. Higher oligomeric states preferentially occupy regions of higher local curvature at WT and moderately weakened LCR-LCR interactions (*ε*_*LCR*−*LCR*_ = 2.8 and 2.1 kcal/mol). This curvature preference diminishes progressively upon further weakening the LCR-LCR interactions: at *ε*_*LCR*−*LCR*_ = 1.4 kcal/mol, the CPSF6 populations become less specific in their tilt distribution, with the multi-oligomer (n = 2) population becoming dominated by a low-tilt mode overlapping n=0 hexamers.

Experiments have demonstrated that abrogating CA-CPSF6 interactions and CPSF6 oligomerization obstructs the capsid at the NPC, preventing passage into the interior of the nucleus. When localized at the nuclear basket, NUP153 also directly engages with the capsid through FG peptides, triple-arginine (RRR) motif, and likely also the prion-like interactions mediated by the LCRs flanking the FG peptide, similar to CPSF6 [26,43-45]. Therefore, seamless handover of the capsid from the nuclear basket to the interior of the nucleus will require CPSF6 oligomerization displacing the pre-assembled NUP153 at the capsid. This handover step is contingent on the enhanced CA binding affinity of CPSF6 relative to NUP153. In particular, the FG peptide of CPSF6 exhibits stronger binding to CA compared to NUP153, which favors the handover. In addition, since LCR-mediated interactions are critical for CPSF6 binding avidity, we hypothesize that LCR_CPSF6-CPSF6_ interactions are energetically dominant over the LCR_CPSF6-NUP153_ and LCR_NUP153-NUP153_ interactions, favoring CPSF6 oligomerization over NUP153 assembly. Therefore, future simulations atomistic resolution (perhaps as backmapped [46] from the present CG ones) will be key for providing a molecular view of the LCR interactions between CPSF6 and NUP153.

## METHODS

### CG modeling and simulation

The details of the CG model development and details of the CPSF6 and capsid model are described in *SI Methods* (S1-S3). All CG MD simulations were performed in the LAMMPS software [47]. The Visual Molecular Dynamics (VMD) software was used to visualize the CG MD trajectories and make snapshots [48].

For the CG MD simulations of CPSF6 binding to the cone-shaped capsid, a cubic simulation box of dimension 300 nm in *x, y*, and *z* directions is used. In the simulation box, the capsid is placed such that the geometric center of the capsid aligns with the center of the box. 600 copies of CPSF6 were initially distributed in the void space in an equispaced grid. The grid to position the CPSF6 copies was created such that each grid point was at least 2 nm from the capsid surface. The system was then evolved for 5 × 10^6^ *τ*_*CG*_ to randomize the initial distribution of CPSF6 proteins. In these simulations, all the CPSF6-CPSF6 and CA-CPSF6 associative interactions were turned off. The simulations were then further evolved for 3 × 10^6^ *τ*_*CG*_, and then configurations saved every 1 × 10^6^ *τ*_*CG*_. These 3 configurations with different spatial distribution of unbound CPSF6 was used as the initial configurations for the CPSF6 binding and oligomerization simulations to the cone-shaped capsid.

The CG MD simulations were performed with a timestep (*τ*_*CG*_) of 50 fs; however, it is again important to note that CG time is not the same as real time, so a 50 fs CG timestep is actually equivalent to a much longer time in terms of the effective sampling achieved by the lower resolution CG model [42]. The equations of motion were integrated with the Velocity Verlet algorithm. The simulations were performed at 310 K in the constant NVT ensemble. The temperature of the CG MD simulations was maintained using the Langevin thermostat with a coupling constant of 1000*τ*_*CG*_ [39]. For analysis and visualization, the simulation trajectory snapshots were saved every 25000 *τ*_*CG*_.

### Analysis

Clustering analysis to identify oligomers was performed on the bound CPSF6 proteins. First, for the bound CPSF6 proteins, we calculated the center of mass of the FG peptide. Then we identify CPSF6 *ij* pairs that are adjacent to each other if the distance between the center of mass of the FG peptide is less than 5 nm. Note that in our CG model, two CPSF6 chains can make a maximum of 32 interchain LCR-LCR contacts. For the adjoining *ij* pairs, we calculated the number of interchain LCR-LCR contacts (the contact is defined if the CG sites are within 2 nm). These adjoining *ij* pairs are classified as part of a cluster if there are at least 16 LCR-LCR contacts (50% of all possible contacts). Oligomers are defined as any cluster with at least three CPSF6 monomers.

For the LCR-LCR contact parameter (*c*_*ij*_) calculation, we calculated all LCR-LCR contacts (both intra- and inter-chain) contacts within CPSF6 oligomers for all the oligomers bound to the capsid. The reported *c*_*ij*_ is the cumulative LCR-LCR contacts (intra and inter) normalized per hexamer (209 for the cone-shaped capsid).

Details of analysis of CA-CPSF6 binding are described in the *SI Methods* (S3).

Details of curvature dependent CPSF6 binding are also provided in the *SI Methods* (S4).

## DATA AVAILABILITY

Model parameters, simulation input files, and custom code will be made publicly available on GitHub/Zenodo upon publication.

## AUTHOR CONTRIBUTIONS

A. H., K. G., and G. A. V. designed the research. A. H. and K. G. built systems, performed simulations, analyzed data, created figures, and prepared the manuscript. All authors contributed to the writing and editing of the final manuscript.

## FUNDING

Research reported in this publication was supported by the Behavior of HIV in Viral Environments (B-HIVE) Center of the National Institute of Allergy and Infectious Diseases (NIAID) of the National Institutes of Health (NIH) under award number U54AI170855. The content is solely the responsibility of the authors and does not necessarily represent the official views of the National Institutes of Health. Computational resources were provided by the Texas Advanced Computing Center (TACC) at The University of Texas at Austin and the Research Computing Center (RCC) at The University of Chicago. Simulations were performed using resources provided by the Advanced Cyberinfrastructure Coordination Ecosystem: Services & Support (ACCESS) program (after September 2022), which is supported by National Science Foundation grants numbers 2138259, 2138286, 2138307, 2137603, and 2138296, and Frontera (at TACC) funded by the NSF (OAC-1818253).

## Notes

### Competing Interest Statement

The authors have declared no competing interest.

