## Supplementary Information for "Mechanistic Insights into HIV-1 Capsid Interactions with CPSF6"

### Supporting Information Methods

#### S1. *CG CPSF6 model*

CG model development of CPSF6 was performed in sequential steps. All-atom (AA) MD simulation trajectories (3 replicas) of three CPSF6 monomers (residues: 261-358) bound to 4 CA hexamers from our previous study were used as a reference from which the CG model of CPSF6 was constructed [1]. The initial configuration for the AA MD simulation was generated using an integrative modeling approach from the X-ray crystal structure of FG peptide bound to CA (PDB ID: 7SNQ), cryo-EM map of $CPSF6_{261-358}$ monomers bound to CA, and sequence-based protein structure modeling [1,2]. The cumulative atomistic statistics (3 replica simulations) were used as the reference to derive the CG molecular interactions. In the AA simulations, each CPSF6 at the N-terminal is capped by a GST dimer which connects the residue 261 with a 12-residue linker. First, we mapped the CPSF6 monomers ($CPSF6_{261-358}$) from the reference AA trajectories. A linear mapping function was used, in which the center of mass of the $C_\alpha$ atoms of three consecutive residues was mapped to a single CG site. The mapped CG $CPSF6_{261-358}$ consisted of 33 sites. Each CPSF6 monomer is modeled as a linear polymer of CG beads where each bead is connected to consecutive beads with flexible harmonic bonds (harmonic bond force constant, $k_{bond}$ = 0.5 kcal mol$^{-1}$ Å$^{-2}$, and equilibrium bond distance, $r_{eq}$ = 8.5 Å). Next, we appended a 20-bead linear polymer to the N-terminal bead of CG $CPSF6_{261-358}$ to model the N-terminal non-LCR segment ($CPSF6_{201-260}$). In the AA MD simulations, the GST dimers act as a crowder to prevent over-aggregation of the LCR segments. Therefore, the non-LCR segment in our CG model implicitly emulates the crowding effect of the GST dimers in AA MD simulations. The AA MD trajectories of 4 CA hexamers and 3 CPSF6 complex was then mapped to the CG representation. The mapping function for CG CA monomers was generated using the Essential Dynamics Coarse Graining (EDCG) protocol, and the details of CG model generation of CA are described previously [3,4].

The CG molecular interactions involving all the protein components and drug molecules are modeled using pairwise repulsive excluded volume ($E_{excl}$) and attractive ($E_{attr}$) interactions. The repulsive excluded volume interaction is modeled with a soft cosine potential, $A\left(1+\cos\left(\frac{\pi r_{ij}}{r_c}\right)\right)$. Here, $r_{ij}$ is the pairwise distance between CG site types $i$ and $j$. The distance cutoff ($r_c$) for the excluded volume interactions between CA-CPSF6 and CPSF6-CPSF6 was set at 1.25 nm. The value of the coefficient $A$ was set at 25 kcal mol$^{-1}$. The pairwise attractive interactions ($E_{attr}$) between the CA and FG peptide of CPSF6 ($\varepsilon_{CA-FG}$) are modeled with the Gaussian potential ($E_{Gauss}$), $\frac{H_{ij}}{\sigma_{ij}\sqrt{2\pi}}\exp\left(-\frac{\left(r_{ij}-r_{0,ij}\right)^2}{2\sigma_{ij}^2}\right)$. Here, $r_{0,ij}$ and $\sigma_{ij}$ are the mean and standard deviation of the distance between CG site types $i$ and $j$. The coefficient $H_{ij}$ is the energy scale for the attractive interaction for a specific $ij$ pair. The $r_{0,ij}$ values for $ij$ pairs were calculated from the CG-mapped AA MD trajectory. $\sigma_{ij}$ value of 0.12 nm was used for all $ij$ pair interactions between CA and FG peptide. The radial cutoff was set at 2.5 nm. The coefficient $H_{ij}$ for the CA-FG $ij$ pairs were optimized using relative-entropy minimization (REM) [5]. In the REM protocol, an initial $H_{ij}$ value was chosen and then iterated using Newton-Raphson method. The REM optimization resulted in a $H_{ij}$ value of -0.85 nm kcal/mol for $\varepsilon_{CA-FG}$.

The CPSF6-CPSF6 associative interactions between the LCR region ($\varepsilon_{LCR-LCR}$) were modeled using a 12-6 Lennard Jones potential ($E_{scLJ}$) with a modified soft-core [6], $4\varepsilon_{LCR-LCR}\lambda^n\left[1/\left(\alpha_{LJ}(1-\lambda)^2+(r/\sigma)^6\right)^2-1/\left(\alpha_{LJ}(1-\lambda)^2+(r/\sigma)^6\right)\right]$. Here, $n$ = 2, $\alpha_{LJ}$ = 0.5, $\lambda$ = 0.6 and $\alpha$ = 1.5 nm. The radial cutoff was set at 3 nm. First, from the cumulative CG-mapped AA

MD trajectory, we determined the CG bead pairs of the LCR segment that are in contact in at least 50% of the total simulation time. We defined a successful contact if the distance between CG beads $i$ and $j$ is less than 2 nm. Intra-chain contacts are only considered if $|i-j|>3$. We then choose the value of the coefficient ($\varepsilon_{LCR-LCR}$) for CG LCR-LCR interactions which reproduces the probability distribution of the cumulative contact parameter ($c_{ij}$) of the CG-mapped AA MD trajectories (**Fig. S4**). For the wild-type CPSF6 we use $\varepsilon_{LCR-LCR}$ = 2.8 kcal/mol. The polymer chain emulating the N-terminal non-LCR segment do not have any associative interactions in our CG model. The mutant CPSF6 model emulating the chimeric proteins lacking the LCR-LCR interactions was modeled by applying a scaling factor to the wild-type $\varepsilon_{LCR-LCR}$. Specifically, we simulated mutant CPSF6 models with $\varepsilon_{LCR-LCR}$ = 1.4 and 2.1 kcal/mol.

### S2. *CG Capsid model*

The CG model of CA monomer consists of 46 sites (~5 residues per CG site). The details of the CG model development of CA are described previously [4]. The intramonomer bonding topology of the CA monomer is modeled as a heterogeneous elastic network with a distance cutoff of 3 nm. The non-bonded CA-CA associative interactions are modeled with the Gaussian potential ($E_{Gauss}$). The radial cutoff was set at 2.5 nm.

### S3. *CA-CPSF6 binding analysis*

To determine if CPSF6 is bound to a monomer in the CG MD simulations, we used a distance metric ($d_{CA-FG}$) between the CPSF6 FG peptide and CA, $\left|\frac{1}{N_{CA}}\sum_{i=1}^{N} r_{CA(i)} - \frac{1}{N_{FG}}\sum_{i=1}^{N} r_{FG(i)}\right|$. Here, $d_{CA-FG}$ is the distance between the center of mass of the selected CG sites of CA and the center of mass of the FG peptide. The FG peptide consists of 5 CG beads. $r$ is the coordinate of the selected CG sites of CA and FG. For CA, CG beads 8 (residue: 34-40), 10-13 (residue: 48-76), 21-22 (all-atom residue: 99-105), 34-37 (residue: 170-182) were used. CPSF6 is classified as bound if $d_{CA-FG}$ < 3 nm.

### S4. *Curvature dependent CPSF6 binding analysis*

To characterize the local curvature of a given hexamer, we developed a scheme to compute the tilt angles between neighboring hexamers. For each hexamer, a best-fit plane encompassing all constituent CG beads was determined using singular value decomposition (SVD). The unit normal vector to this best-fit plane was then computed. The acute angle between the unit normal vectors of two neighboring hexamers was defined as the corresponding hexamer-hexamer tilt angle, given by $\theta_{ij} = \arccos\left(\left|\hat{\mathbf{n}}_i \cdot \hat{\mathbf{n}}_j\right|\right)$. The local curvature of each hexamer was quantified as the mean of the tilt angles between that hexamer and each of its six nearest neighboring hexamers (**Fig. 4** and **Fig. S5**). A CPSF6 monomer is considered bound to a given hexamer if the distance between its FG-motif centroid and any monomer of that hexamer is less than 3 nm. For each pair of bound CPSF6 monomers, the distance between their FG-motif centroids is then computed. Pairs separated by less than 7.5 nm are considered connected. CPSF6 oligomers are subsequently identified by constructing connected chains of CPSF6 monomers, where each consecutive pair of monomers is connected by an FG-centroid distance of less than 7.5 nm. Any connected chain containing three or more CPSF6 monomers is classified as a CPSF6 oligomer. Since every monomer within an identified oligomer is bound to a hexamer, each oligomer is assigned to the hexamer to which the majority of its constituent monomers are bound. The distributions shown in **Fig. 4** of the main text correspond to the resulting oligomer count as a function of the local hexamer tilt angle.

### Supporting Information Figures

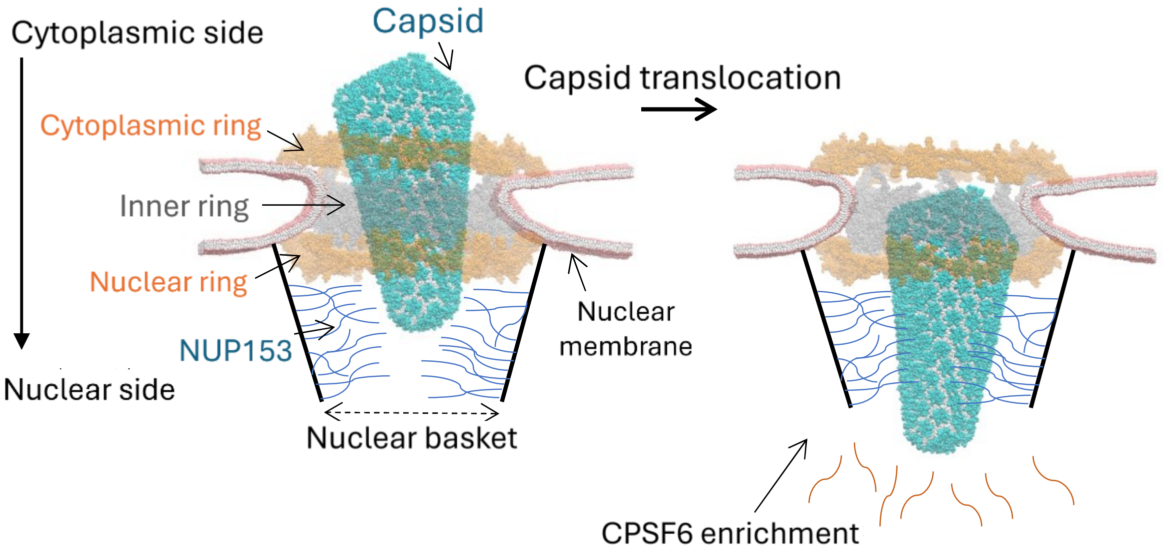


**Fig. S1** Model of the HIV-1 capsid nuclear entry. The $CA_{NTD}$ and $CA_{CTD}$ domain of the HIV-1 capsid is shown in cyan and white beads respectively. The cytoplasmic and nuclear ring of the nuclear pore complex (NPC) is shown in transparent orange beads. The inner ring of the NPC is shown in transparent silver beads. The nuclear membrane is shown in pink and white beads. A schematic of the nuclear basket is added to the NPC model. The intrinsically disordered NUP153 and CPSF6 are shown in blue and orange curved lines. Note, the capsid translocation from the cytoplasmic side to the nuclear side of the NPC.

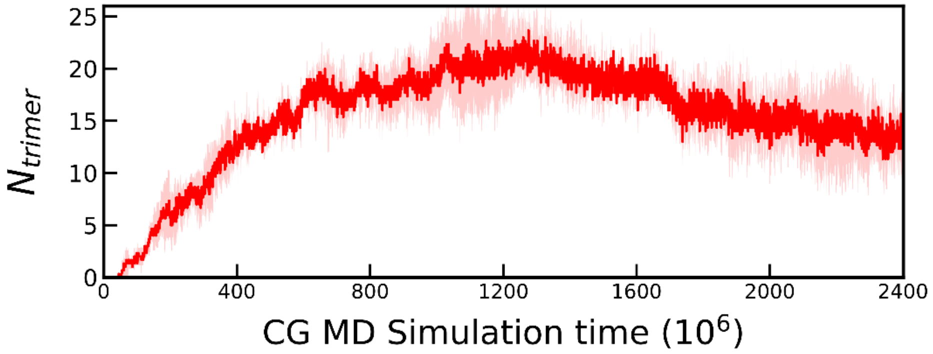


**Fig. S2** The time series plot of total WT CPSF6 trimers ($N_{trimer}$) bound to the capsid during the oligomerization simulations. The mean and standard deviation values are calculated from the endpoint of the simulation trajectories for 3 replicas.

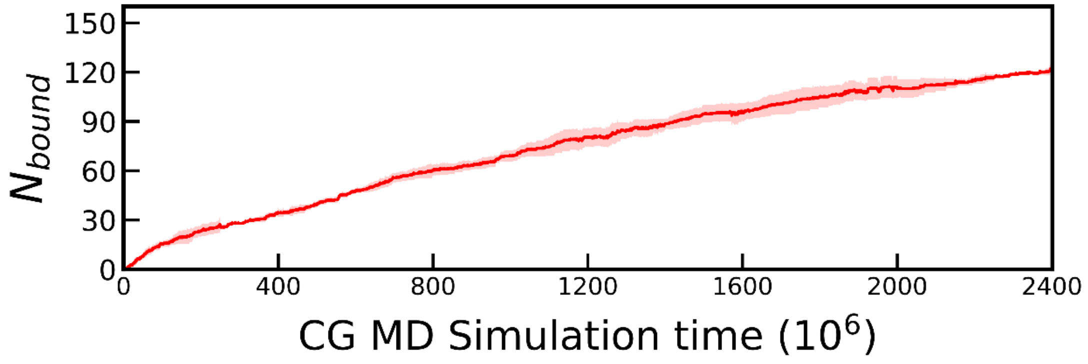


**Fig. S3** Mutated CPSF6 binding to the cone-shaped capsid. The time series plots of total mutated CPSF6 ($\varepsilon_{LCR-LCR}$ = 0.0 kcal/mol) bound to the capsid lattice ($N_{bound}$). The mean and standard deviation values are calculated from the endpoint of the simulation trajectories for 3 replicas.

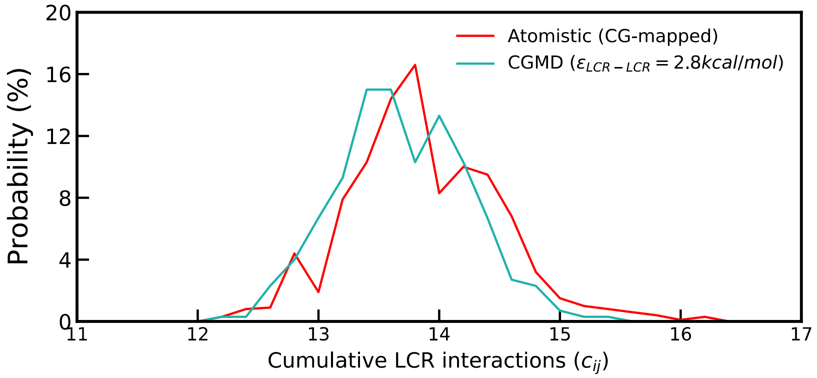


**Fig. S4** Probability distribution of the cumulative LCR-LCR contact parameter ($c_{ij}$) for the CG-mapped atomistic trajectory (red line) and CGMD simulation (green) for $\varepsilon_{LCR-LCR}$ = 2.8 kcal/mol. The calculations were performed for 3 CPSF6 chains bound to 4 CA hexamer lattice (**Fig. 1** in the main text).

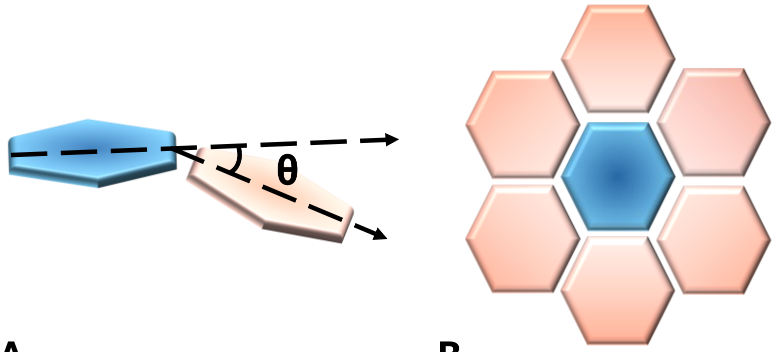


A B

**Fig. S5** Illustration of the hexamer-hexamer tilt angle. **(A)** Two neighboring hexamers and the tilt angle between them. In our calculations, the tilt angle is defined as the angle between the normal vectors to the best-fit planes of the two hexamers. **(B)** A 'central' hexamer and its six nearest neighboring hexamers. The local tilt of the central hexamer is defined as the mean of the tilt angles between the central hexamer and each of its six neighbors, which serves as a measure of its local curvature.

**Supporting Information References:**
